# Analyzing cannabinoid-induced abnormal behavior in a zebrafish model

**DOI:** 10.1101/2020.07.13.200428

**Authors:** Akihiro Hasumi, Hideyuki Maeda, Ken-ichi Yoshida

**Affiliations:** Department of Forensic Medicine, Tokyo Medical University, Tokyo, Japan

**Keywords:** repeated light and dark test, cannabidiol, WIN55,212-2, zebrafish, marijuana

## Abstract

This study investigated the locomotor activity and responses under stressful conditions to assess marijuana-induced abnormal behavior in zebrafish larvae (*Danio rerio*), as an alternative to the standard rodent model. To induce the desired symptoms, cannabidiol and WIN55,212-2, two chemical components derived from marijuana plants, were used. A repeated light and dark test was used to assess how drug exposure influenced locomotive responses. Subjects were examined after moderate cannabidiol and WIN55,212-2 exposure and 24 h after relocation into untreated water. Cannabidiol did not produce a dose-dependent inhibitory effect on locomotor activity; 0.5 and 10 μg/mL cannabidiol decreased total distance and velocity, and 10 μg/mL cannabidiol attenuated responses in the darkness. No differences between the control and cannabidiol-treated groups were apparent after 24 h in fresh water. WIN55,212-2 at 0.5 and 1 μg/mL almost completely stopped fish activity even in darkness and at 10 μg/mL induced mortality. Spending 24 h in fresh water almost reversed drug-induced immobilization, even in WIN55,212-2-treated groups. Subjects were evaluated for responses to additional cannabidiol after WIN55,212-2 exposure. Cannabidiol attenuated WIN55,212-2-induced abnormal immobilization, whereas commensurate doses of cannabidiol and WIN55,212-2 produced a mixed response. In conclusion, the behavioral effects of marijuana depend on the ratio of the two different cannabinoid properties. The repeated light and dark test might be a suitable method for drug-induced behavioral assay.

## Introduction

Marijuana, also known as cannabis, is an illegal drug derived from *Cannabis sativa* plants that can directly induce psychotropic effects with typical symptoms, such as euphoria and addiction [1, 2]. Habitual users of marijuana or marijuana-like drugs are likely to be involved in fatal driving accidents [3, 4], and suffer from psychopathic behaviors, and schizophrenia-like psychosis [5]. In addition, abuse of marijuana or similar psychotropic drugs can induce serious adverse effects, including sudden death [6, 7]. Some case reports have even suggested that several overdoses on these drugs may increase the risk of serious cardiac conditions, including arrhythmia, myocardial infarction, cardiomyopathy, and death from cardiac arrest [8]. Thus, in fatal accidents involving users with marijuana who had exhibited abnormal behavioral symptoms, autopsies should attempt to determine whether the cause of death was an illness drug-induced behavioral or physical dysfunction, or trauma from the crash.

Given the potential dangers, the drug components that cause these side effects need to be identified. Several assays have been developed that evaluate locomotor activities, startle latency, behavior modifications, and physiological conditions including cardiac arrest [9–11]. Typically, rodents are the preferred model for such studies owing to their reliability in behavioral screenings, and physiological responses [12–14]. However, during the past decade, zebrafish (*Danio rerio*) have become increasingly popular as a model for pharmacological or behavioral research testing diverse psychotropic drugs [15–19]. Zebrafish are easy to handle and are more cost-effective than rats. Furthermore, zebrafish have some genetic phenotypes, and specific proteins that are similar to humans [20].

Locomotion in zebrafish is usually evaluated using three variables: moved distance, moving velocity, and moving duration [21–24]. However, these three items are insufficient evidence of abnormal behaviors (including drug-induced symptoms). When attempting to duplicate pharmacological symptoms, researchers can also concurrently measure the sensitivity of responses or startle latency using a light or dark preference test [25]. Analyses of the residual effect on behavioral functions after the withdrawal of the drugs, and the interaction effects should be added to the study [26,27]. In both rodent and zebrafish models, typical locomotor activity can be examined using moved distance, moving velocity, and moving duration. Only rodents, however, can participate in blood examination to evaluate drug concentrations [28]. Therefore, researchers interested in zebrafish as a pharmacological model need to identify appropriate alternatives.

In this study, the repeated light and dark test for stress sensitivity screening in addition to typical locomotor activity was selected. Cannabidiol (CBD) and R-(+)-[2,3-Dihydro-5-methyl-3-[(morpholinyl)methyl]pyrrolo[1,2.3-de]-1,4-benzoxazinyl]-(1-naphthalenyl)methanone mesylate (WIN55,212-2; WIN)—two typical cannabinoids—were used to induce marijuana-like behavior. The pharmacological effects of WIN are very similar to delta-(9)-tetrahydrocannabinol (THC), which is derived directly from marijuana and activates cannabidiol 1 and 2 receptors [29,30]. Beyond those findings, no studies are available that investigate whether CBD-treated groups fully return to their pre-WIN-treated state in zebrafish.

The objective of this study was to determine how CBD interacts with the abnormal behaviors induced by WIN. We hypothesized that CBD would attenuate WIN-induced abnormal behaviors. We evaluated zebrafish larvae locomotion during acute drug exposure and drug withdrawal, as well as whether repeated light or dark stimuli, as a measure of stress sensitivity, would affect drug-induced locomotive responses. Stress sensitivity tests are common in the rodent model. The repeated light and dark test was duplicated after drug-treated zebrafish were placed in drug-free water for 24 h. Interactional effects on activity were assessed by adding CBD after exposure to different WIN concentrations. This research is particularly relevant in the field of forensic toxicology, where the toxic effects of interactions between CBD and WIN can result in sudden deaths. These results will elucidate potential links between cannabinoids and unexpected abnormal behaviors that could trigger fatal accidents or physical dysfunctions.

## Materials and Methods

This study employed the repeated light and dark test to assess drug-induced activity and stress sensitivity because fish can perform the necessary movements during repeated, alternating periods of alert action (light-on), or escape action (light-off). Typically, animal trials investigating pharmacological effects are conducted immediately after drug administration, or after replacing the drug solution with fresh water. We included an extra assessment of drug-treated fish which were housed for 24 h in fresh, unmedicated water. The study used a high-throughput tracking system (Danio Vision XT) and behavioral analysis software (Ethovision XT 11.5) [31], which is currently rare among researchers using small organisms but is becoming increasingly common for investigating pharmacological effects in zebrafish.

### Fish and fish culture

The Experimental Animal Committee of Tokyo Medical University approved all experiments (approval number: H30-0020, R1-0119). Healthy, adult zebrafish (*Danio Rerio*; wild-type, purchased from Kamihata Fish Industries Ltd., Tokyo, Japan) were housed and raised in aerated breeding units at a density of 10 fish per liter, in water from a recirculating water system supplied with dechlorinated municipal tap water. They were fed flake food twice daily. Fish were kept in conditions of pH 7.5–8.0, conductivity of 300–500 μS/cm, a temperature of 26–28 °C, and a 14:10 h light:dark photoperiod. To produce embryos, male and female zebrafish were paired in the evening and fertilized embryos were collected from the mated zebrafish and placed in petri dishes with fresh water. Embryos were transferred to a 28 °C incubator on a 14:10-h light:dark cycle until 4–6 days post fertilization (4–6 dpf). Exposed embryos were screened to assess overall health and dead embryos were removed daily.

### Chemicals

CBD was diluted with 0.05% methanol, and sterile saline (Cayman Chemical, Ann Arbor, MI). CBD, one of the major compounds present in the marijuana (*C. sativa*) plant, has some medicinal properties, however, its mechanism is not well known [32, 33]. WIN (Adooq Bioscience, Irvine, CA) was diluted with 0.05% dimethyl sulfoxide (Dojindo Laboratories, Japan) and sterile saline. WIN is a synthetic cannabimimetic agent with a high affinity for cannabinoid 1 and 2 receptors, two mixed cannabinoid receptor agonists like THC [34]. WIN accounts for most of the effects elicited by marijuana [35]. Thus, WIN is an alternative drug to THC, the traditional psychotropic compound derived from marijuana plants and is used primarily to duplicate pharmacological symptoms of marijuana by researchers. Both drugs (THC and WIN) indicate mild enhancement of the brain-stimulation reward [36].

### Experimental procedure

Zebrafish larvae (4–5 dpf) were maintained in 96-well microtiter plates (1 larva/well) (IWAKI Co., Ltd. Tokyo, Japan) filled with 300 μL E3 medium (n = 96). Locomotor activity was assessed after CBD and WIN treatments of 0.5, 1, 5, and 10 μg/mL (16 larvae/group) were added directly into the water in the well. CBD+WIN treatments of 0.5+0.5, 1+0.5, 5+0.5, 10+0.5 μg/mL (16 larvae/group), and CBD+WIN treatments of 0.5+1, 1+1, 5+1, 10+1 μg/mL (16 larvae/group) were also assessed (n = 384). Fish in each treated group were then subjected to repeated light and dark tests to determine the differences in locomotor activities and responses, examined under the alternate light or dark circumstances at 15 minutes intervals. Drug effects on subjects were measured for 3 h using Danio Vision XT and Ethovision XT 11.5 (Noldus Information Technology, Wageningen, The Netherlands). Vision is a high-throughput system designed for studying locomotion in small organisms and can simultaneously track up to 96 individuals. This experiment examined ambulation, total distance traveled, velocity of movement, and moving duration. These parameters were measured again 24 h after the subjects were placed into fresh, untreated water during drug withdrawal. We omitted the results of reactions in treated groups, when more than half the fish in the groups died beforehand.

We conducted pre-experimental trials and gauged that zebrafish larvae displayed a startle action for a few seconds following sudden light stimulus, and instantly followed this by moderate movements in the light, whereas zebrafish larvae displayed panic and escape actions for 5–15 min after a sudden dark stimulus, which was gradually followed by decreasing movements in the dark. The results of repeated light stimuli were likely to diverge at the first and second test times, whereas the results from the third to sixth time converged and corresponded to those of the control fish. However, all the results that follow may be attenuated gradually because of fatigue or acclimation. Our previous study suggests that the interval between light or dark states should be within 15 min of measurement to avoid acclimation of reactions to those stimuli.

### Statistical analysis

Data are presented as means ± SEMs (standard error of the mean). Statistical analysis was performed in GraphPad Prism 6 for Windows version 6.05. The data were analyzed using Dunnett’s test, the Tukey–Kramer test, or Student’s t-test at a confidence level of 95%.

## Results

### Effects of CBD at different doses during drug exposure and 24-h drug withdrawal

CBD at a dose of 10 μg/mL significantly influenced locomotor activity (P < 0.05; Fig 1. A), and induced decreases in total distance and velocity. CBD at doses of 5 and 10 μg/mL induced increases in locomotor activity compared with the dose of 0.5 μg/mL. After 24 h without the drug, however, the differences were not significant compared with the control (Fig 1. B). CBD at a dose of 10 μg/mL exerted the strongest effect (low responses to darkness) compared with control (P < 0.01; Fig 2. A), and the lowest responses occurred at intervals three, five, and six. The differences in locomotor responses were not significant for the remaining drug-treated groups. Locomotor responses recovered even at 10 μg/mL, after 24 h withdrawal (Fig 2. B). In the acute trials, the responses tended to decrease both in the light and the dark without significant differences except at 10 μg/mL. After 24-h withdrawal, the responses of fish receiving 0.5 and 1 μg/mL had almost recovered (Fig 2; B). Responses at doses of 5 and 10 μg/mL increased overall without significant differences (Fig 2; B).

**Fig 1.**
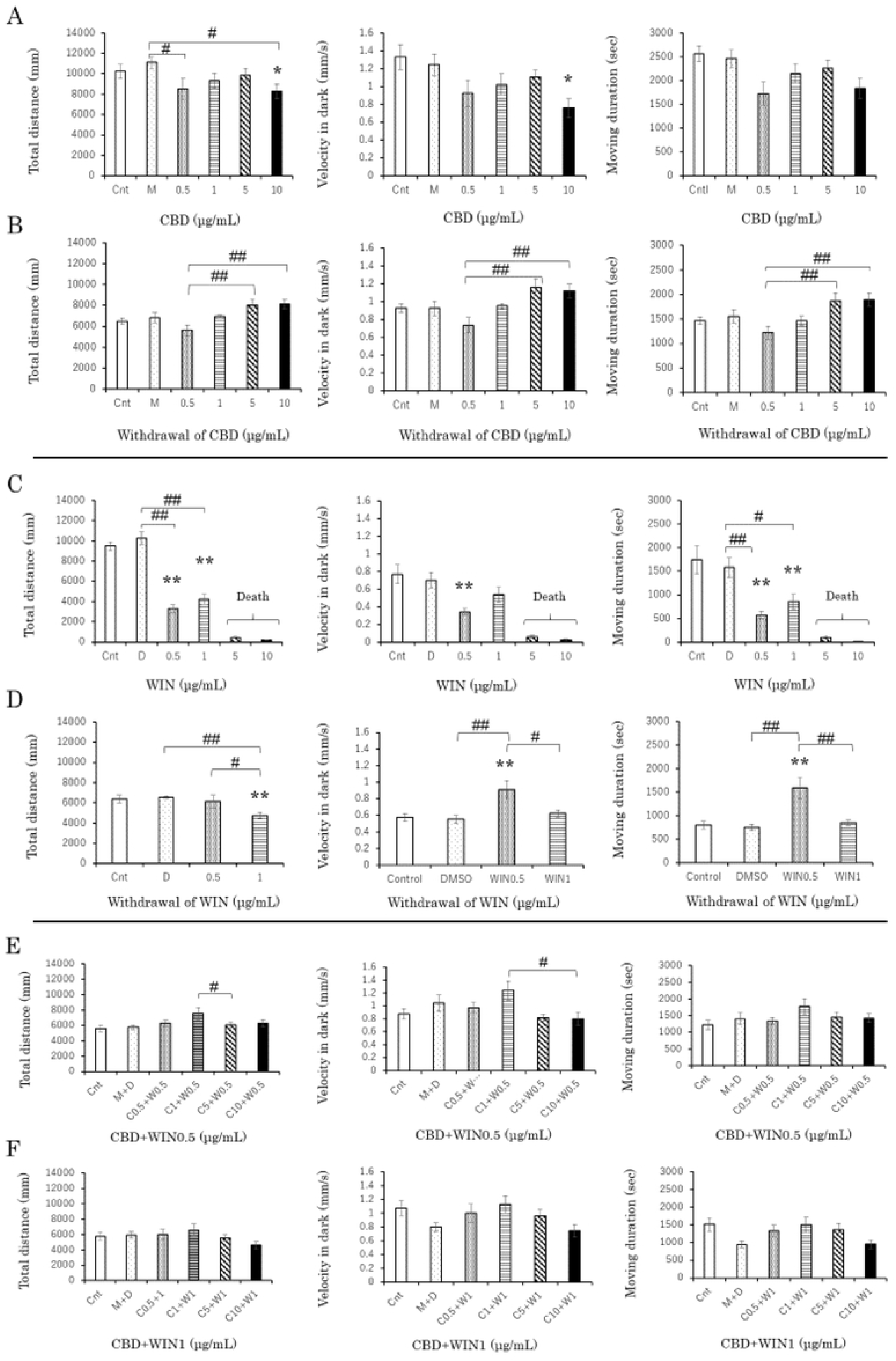
Locomotor effects of cannabidiol (CBD), WIN55,212-2 (WIN), and CBD+WIN at different doses (each experiment; n = 96). CBD (10 μg/mL) significantly induced decreases (A). CBD (5, 10 μg/mL) induced increases compared with a dose of 0.5 μg/mL (B) after 24 h withdrawal. WIN (0.5, 1 μg/mL) significantly induced decreases (C). WIN (0.5 μg/mL) induced increases (D) after 24-h withdrawal. CBD1 + WIN0.5 induced significant differences compared with CBD5 + WIN0.5 (E), and CBD10 + WIN0.5 (E). Cnt; control, M; methanol, D; dimethyl sulfoxide. *P < 0.05, **P < 0.01 vs control, #P < 0.05, ##P < 0.01 vs drug-treatment. Tukey’s multiple comparison test. All values are expressed as mean ± SEM.

**Fig 2.**
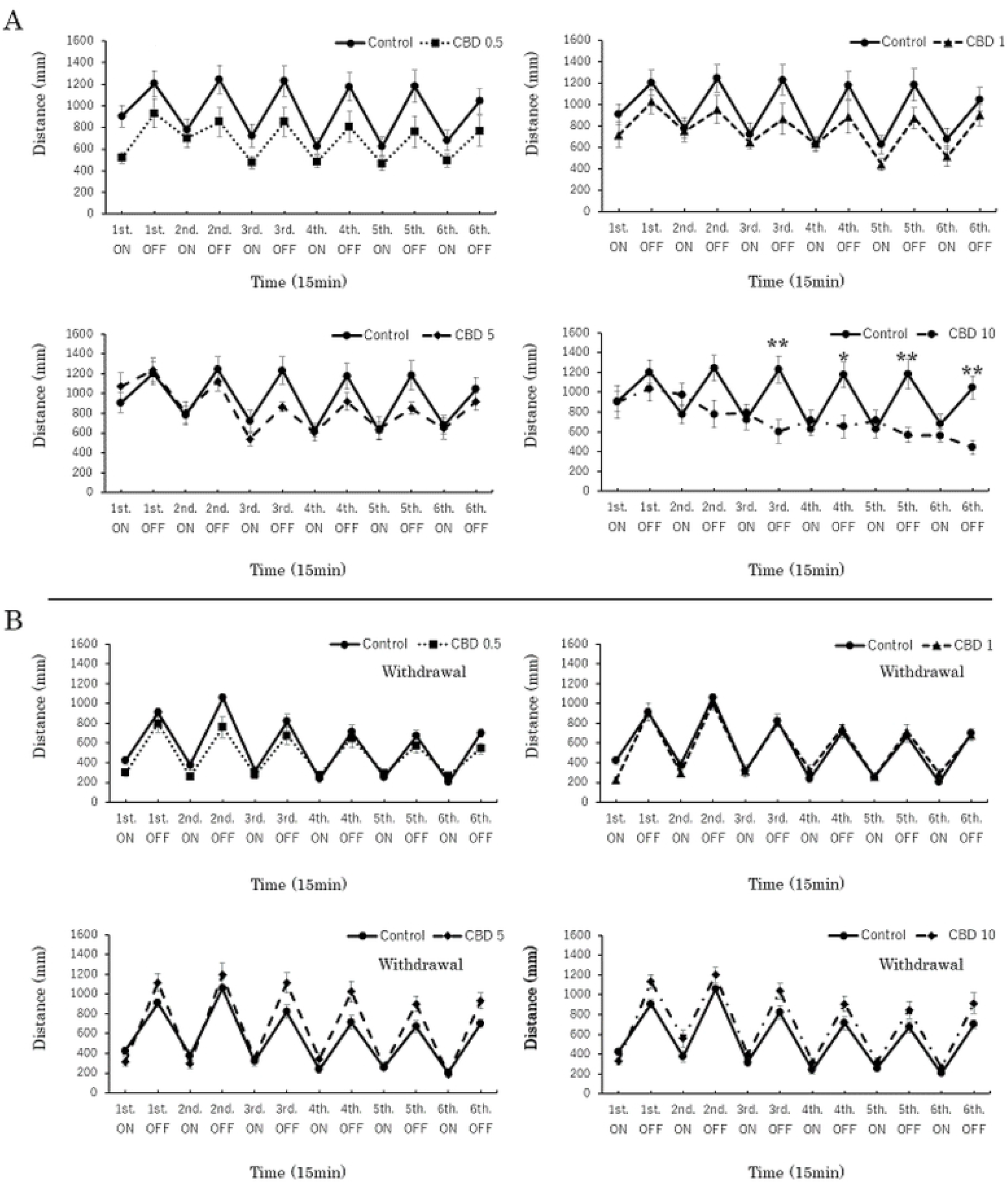
Responses to cannabidiol (CBD) at different doses during drug exposure and 24-h withdrawal (each experiment; n = 96). CBD (10 μg/mL) exerted the strongest effect (low responses to darkness) compared with the control (A). Locomotor responses recovered at 10 μg/mL after 24-h withdrawal (b-4). *P < 0.05, **P < 0.01 vs control, #P < 0.05, ##P < 0.01 vs drug-treatment. T-test. All values are expressed as mean ± SEM.

### Effects of WIN at different doses during drug exposure and 24-h withdrawal

WIN doses of 0.5, and 1 μg/mL significantly influenced locomotor responses (P < 0.01; Fig 1. C). The two doses of WIN induced decreases in locomotor activity and dulled responses to the repeated light and dark test (Fig 3. A). However, doses of 5 and 10 μg/mL were lethal in less than 24 h (Table 1). Over half of fish in the 5 μg/mL group were dead, and all fish in the 10 μg/mL group died within the 24 h. All fish were still alive at the end of the acute trials. Thus, we omitted the results of both the 5 and 10 μg/mL groups from statistical analysis. Doses of 0.5 and 1 μg/mL induced increases in locomotor activity after 24-h withdrawal (P < 0.01; Fig 1. D). After 24-h withdrawal, the responses in the light recovered at a dose of 0.5 μg/mL. Low activity remained at the first interval in both states, and at the fourth interval in light. An increased response in the sixth interval in the dark was observed (P < 0.01; Fig 3. B). Attenuation of both responses in the first and fourth intervals remained significant, and the 1 μg/mL dose corresponded to the control from the sixth interval (P < 0.01; Fig 3. B).

**Fig 3.**
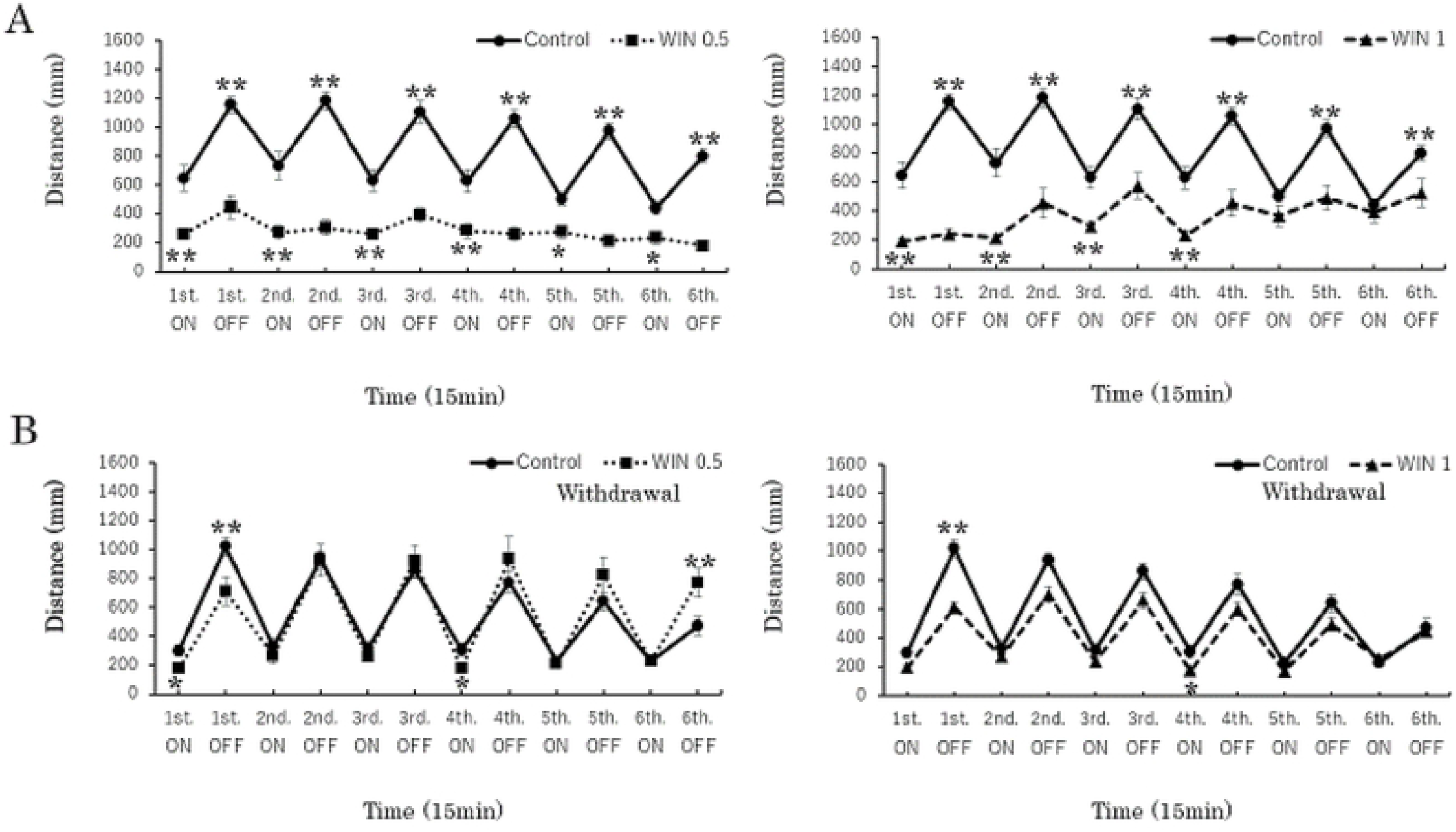
The responses to WIN55,212-2 (WIN) at different doses during drug exposure and 24-h withdrawal (each experiment; n = 96). WIN (0.5, 1 μg/mL) induced significant decreases in the response to both light and dark (A). After 24 h without the drug, at 0.5 μg/mL, there was low activity in the first interval in both states, and in the fourth interval in light. Increasing responses in the sixth interval in dark were seen (B). The 1 μg/mL dose showed that the attenuation of both responses in the first and fourth intervals remained significant (B). *P < 0.05, **P < 0.01 vs control, #P < 0.05, ##P < 0.01 vs drug-treatment. t-test. All values are expressed as mean ± SEM.

**Table 1.** Lethal effects of drugs (each experiment; n = 96).

| Lethal effects of drugs |  |  |  |
| --- | --- | --- | --- |
| Drug | Concentration | Percentage | Lethal number / Total number |
| WIN | 5 (µ g/mL) | 60 % | 10 / n=16 |
|  | 10 (µ g/mL) | 100 % | 16 / n=16 |
| WIN+CBD (W+C) | W5+C5 (µ g/mL) | 50 % | 8 / n=16 |
|  | W10+C10 (µ g/mL) | 100 % | 16 / n=16 |

WIN55,212-2 (WIN) doses of 5 and 10 μg/mL were lethal at < 24 h. Over half the fish receiving 5 μg/mL were dead, and all fish receiving 10 μg/mL died within 24 h. However, all zebrafish were still alive at the end of the acute trial.

### Interactions between CBD and WIN at different doses during drug exposure

Interactions between WIN (0.5 and 1 μg/mL), and CBD (0.5, 1, 5, and 10 μg/mL) changed locomotor responses. CBD doses of 0.5, 1, 5, and 10 μg/mL attenuated the low activity (P < 0.05; Fig 1. E, F), and dull responses induced by the WIN-treated groups (P < 0.01; Fig 4. A, B). Thus, CBD induced increases in locomotor activity, and responses to repeated light and dark states after 0.5 and 1 μg/mL doses of WIN. Responses increased at the fifth and sixth intervals in dark, when the CBD-treatment at a dose of 1 μg/mL (double doses of CBD 0.5 μg/mL) was added to pretreated groups of WIN 0.5 μg/mL (P < 0.01; Fig 4. A). Responses decreased in the second to sixth intervals in the dark when 10 μg/mL CBD was added to pretreated WIN 0.5 μg/mL (P < 0.01; Fig 4. A). In contrast to WIN0.5 + CBD1 (μg/mL), responses decreased in the fifth and sixth intervals in the dark, when 10 μg/mL CBD was added to pretreated 1 μg/mL WIN (P < 0.01; Fig 4. B). The shared effect of CBD and WIN (0.5 and 1 μg/mL) demonstrated that CBD and WIN in 1:1 ratio induced recovery (Fig 4. A, B). CBD5 + WIN0.5 (μg/mL), CBD0.5 + WIN1 (μg/mL), and CBD5 + WIN1 (μg/mL) tended to slow the responses in the first interval in the dark. Significant attenuation was seen in CBD5 + WIN1 (μg/mL) (P < 0.05; Fig 4. A, B).

**Fig. 4.**
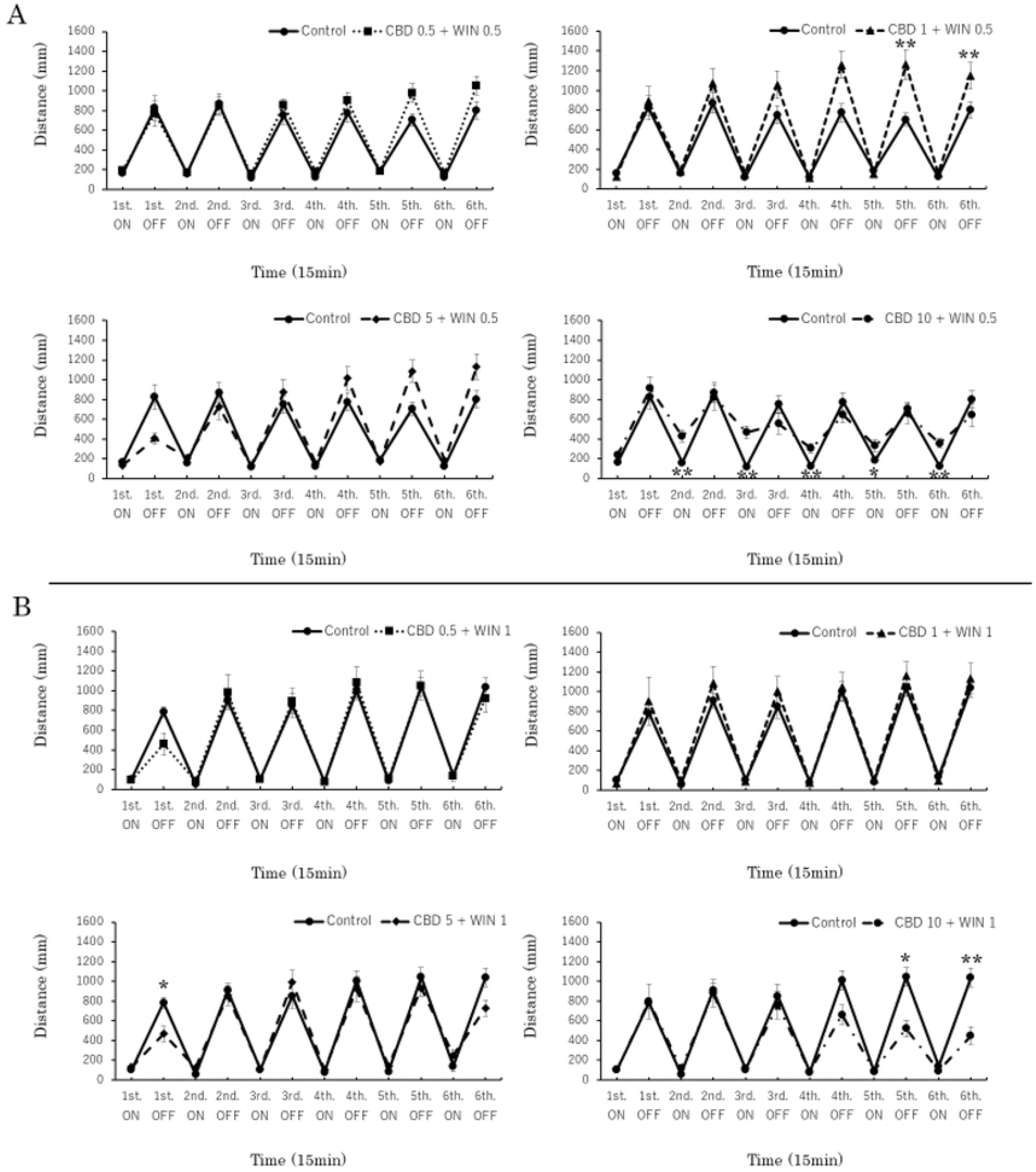
Responses to interactions between cannabidiol (CBD) and WIN55,212-2 (WIN) at different doses during drug exposure (each experiment; n = 96). Interactions between WIN (0.5 and 1 μg/mL) and CBD (0.5, 1, 5, and 10 μg/mL) changed the locomotor responses. Interactions of CBD1 + WIN0.5 (μg/mL) induced significant increases in the fifth and sixth intervals in the dark (A), whereas CBD10 + WIN0.5 (μg/mL) induced decreases from second to sixth intervals in the light (A). Interactions of CBD5 + WIN1 (μg/mL) induced decreases in the first interval in dark (B), whereas CBD10 + WIN1 (μg/mL) induced decreases in the fifth and sixth intervals in the dark (B). *P < 0.05, **P < 0.01 vs control, #P < 0.05, ##P < 0.01 vs drug-treatment. T-test. All values are expressed as mean ± SEM.

## Discussion

Dysfunction of serotonergic neurons that huddle together to form raphe nuclei are clinical characteristics of Parkinson’s disease [37]. The mechanism of serotonergic activation constitutes a substantial portion of the ascending reticular activating system in the brain stem, known as the arousal and conscious center, along with the locus coeruleus [38]. Normally, sunlight stimuli pass through the retina to the suprachiasmatic nucleus, followed by gradual substitution from melamine to serotonin, activating the reset functions in the circadian cycle [39]. Light stimulus was used as a startle stressor because of the hypothesis that serotonin is important in alert or escape actions. Serotonin might be important in rapid response because the adrenal cortical hormone, known as the stress hormone, may exert a delayed action to cope with rapid stressors. Moreover, persistent stresses induce serotonin depletion causing depressive psychosis [40, 41], which is associated with anxiety behavior.

In this study, we hypothesized that moderate startle responses, suggesting alert action, would occur in the light state, and that intense responses like escape action would occur in the dark state. WIN activates cannabinoid 1 and 2 receptors. Even at a lower WIN concentration (0.5 μg/mL), ambulation activity and responses of zebrafish larvae decreased, indicating that these levels are sufficient to inhibit serotonin 5-HT1A receptors, thus activating cannabinoid 1 and 2 receptors. This would eventually alter serotonin secretion and, therefore, behavioral activity. In zebrafish, serotonergic activity is associated with anti-epileptiform activity, suggesting a link between epilepsy and epilepsy-like immobilization, such as catalepsy [42]. Reportedly, activating 5-HT1A receptors via CBD attenuates some distinct drug-induced catalepsy in mice [32, 43]. Serotonin mediated by cannabinoid receptors may also regulate noradrenergic and dopaminergic neurotransmission because cannabinoids (WIN) modulate motoric responses by activating dopaminergic and glutamatergic neurons [44].

Gradual light stimuli induce moderate serotonin elevation, along with secretions of catecholamines including noradrenaline and dopamine, to elevate blood pressure and body temperature for proper activity in circadian rhythmicity [45]. The CBD trial assumed that moderate serotonin induced proper noradrenaline and dopamine elevation, whereas an excess (or surge) of serotonin could inhibit these catecholamines. Startle responses may reflect combinations of the reticular activating system such as the arousal center in the brain stem, basal ganglia as the kinetic coordinating unit, and the limbic system associated with anxiety or fear, which are mediated by the serotonin and catecholamine cascade reaction. Locomotor responses may depend on the initial serotonin neurotransmitters for zebrafish.

Psychoactive drugs, such as WIN or THC, that activate cannabinoid receptors cause hypothermia and hypoactivity (including catalepsy-like immobilization), which are the inverse of the symptoms of serotonin syndrome [46, 47]. Moreover, these drugs also induce euphoria and addiction due to effused dopamine in the nucleus accumbens, such as amphetamines [48, 49]. Hypoactivity or catalepsy-like immobilization could occur because of a shortage in dopamine supply in the basal ganglia. Dopamine is released in the nucleus accumbens [48] and is linked to environmental reward-predictive stimuli [49]. Euphoria and sedative mechanisms may be strongly related to cannabinoid receptors because endogenic cannabinoid (anandaminde) not only activates the cannabinoid receptors, but also the transient receptor potential vanilloid 1 (TRPV1). Cannabinoid links with TRPV1 in regulating some neural functions such as membrane excitability and synaptic transmission [50]. WIN was reported to exert both excitatory and inhibitory effects, neither of which appear to be mediated by cannabinoid 1 and 2, or TRPV1 receptors [51]. Therefore, the pharmacological interaction between cannabinoid and TRPV1 could represent a promising approach for a future strategy to challenge hyperexcitability-based diseases [50]. Although WIN inhibits TRPV1 for capsaicin-evoked responses in nociceptors, and evokes peripheral antihyperalgesia [52], the mechanism for cannabinoid-induced inhibition of TRPV1 in primary sensory neurons is not understood [53].

Suffering under a stressful state or pain continuously activate sympathetic nerves and are followed by activations of vanilloid or cannabinoid receptors for beta-endorphin release [54, 55] to attenuate those stress reactions by means of sedation or even euphoria (“runner’s high”). Serotonin also facilitates beta-endorphin release in the nucleus accumbens [56].

Some studies report that CBD or 5-HT receptor agonists have medical effects such as_antiepileptic seizures [57]. The THC contained in marijuana may have similar effects as endogenic cannabinoids that activate cannabinoid receptors for self-defense mechanisms, whereas the CBD contained in marijuana may have a role in the regulation of other cannabinoids including THC. Previous studies revealed the developmental effects of CBD and THC in zebrafish; however, further studies are needed to assess the latent effects [58]. The proportions of cannabinoids are different because many *Cannabis* plants grow naturally. Thus, the pharmacological effects of marijuana may be hypothesized to be rich in diversity and may depend on the growth area, developmental conditions, circumstances, extraction methods, and processing methods.

The toxic properties of WIN are much higher than those of CBD. Therefore, when it comes to sudden death associated with marijuana, we should recognize that THC or THC-like drugs such as WIN are likely to inhibit sympathetic nerves rapidly and promote cardiopulmonary dysfunction. Both CBD and WIN (even at a low dose) attenuated locomotor activity and responses. WIN demonstrated a stronger principal toxic action than CBD, whereas high CBD doses noticeably attenuated intense escape-like activity in the dark. After 24-h withdrawal, recovery was seen in both pretreated groups. An interaction between WIN and CBD increased activity and even indicated comparative recovery. Reactions varied with different combinations of CBD and WIN doses. A much higher CBD dose than WIN dose, in combination, induced hyperactivity in light periods, indicating that the serotonin-induced reset mechanism for circadian rhythms may be malfunctioning.

In the repeated light and dark test, the results in the first and second intervals may be dispersed, whereas the results from the third interval or later are likely to be more representative of the actual state. Therefore, further study is needed to confirm the behavioral analysis method or manner in zebrafish.

Studying pharmacological effects using marijuana-like drugs in zebrafish is growing in popularity because cannabinoids (such as CBD) are increasingly being used for medicinal purposes worldwide [57, 59]. Our study augments other research on cannabinoid biology, cannabinoid 1 and 2 receptors, and 5-HT1A function in zebrafish [60–62]. Locomotion with light stimuli using zebrafish has been previously evaluated [22, 60, 63, 64]. The results of the current study showed that light attenuated activity, while darkness facilitated activity in normal zebrafish.

In summary, the repeated light and dark test is an appropriate substitute for the startle latency test for evaluating the responses of zebrafish under stressful conditions. CBD and WIN temporarily induced locomotive disorders. Drug withdrawal for 24 h attenuated drug-induced low activity. It was, therefore, concluded that measuring symptoms during and after drug exposure is a valid method for investigating pharmacological effects in a fish model. Our study has important implications for the persistence of drug complications.

The low toxicity of marijuana makes it less important to forensic medicine than other psychomimetic drugs because it is rarely linked to sudden deaths. However, some research suggests that marijuana dangers may be underestimated. For example, catalepsy can also be induced by THC [12]. Although catalepsy is a typical symptom that represents psychosis-induced immobilization and is not recognized as a critical illness associated with sudden death, it can induce serious accidents including fatal car crashes. Like Parkinsonism, catalepsy is likely caused by the disturbance of dopamine 2 receptors in the corpus striatum [14]. In this case, drug-induced catalepsy-like immobilization could be the cause of driving dysfunction if there is evidence linking fatal car accidents with a history of psychotropic cannabinoid (e.g., marijuana) use. However, if fatal traffic accidents are only associated with cannabinoid receptor-agonist intakes such as WIN or THC, then driving dysfunctions would typically be attributed to cannabinoid receptor-induced immobilization. Our study suggests that other cannabinoids, like CBD, could actually cause the abnormal behavior together with the enhanced abnormal behaviors from marijuana intake, which could then combine to result in various symptoms. Thus, we recommend that forensic scientists perform inquests into the causes of marijuana-related deaths until the presence or absence of any correlation between the THC concentration and the contents of other cannabinoids such as CBD is firmly established. The results of our study suggest that forensic screening for marijuana-induced abnormal behaviors in fatal accidents should advance considerably to become useful in judicial cases.

Future studies using zebrafish models should determine whether CBD can attenuate other cannabinoid-induced abnormal behaviors or side effects from psychomimetic drugs, including antipsychotic agents. Several studies have attempted to duplicate and assess marijuana-induced motor functional disorders in both zebrafish and rodents but have had little success in accurately replicating results from rodent models. Thus, further research is necessary to evaluate toxic effects or drug-induced abnormal behavior in zebrafish. This research is particularly relevant in the field of forensic toxicology, where the toxic effects of interactions between CBD and WIN can result in sudden deaths. Our results will help elucidate potential links between cannabinoids and unexpected abnormal behaviors that could trigger fatal accidents or physical dysfunction.

## Acknowledgments

This study was supported by the Department of Forensic Medicine at Tokyo Medical University, and the Department of Pathophysiology at Tokyo Medical University. We gratefully acknowledge their support.

We would like to thank Editage (www.editage.jp) for English language editing.

